## Supplementary material for "Simultaneous selection of nanobodies for accessible epitopes on immune cells in the tumor microenvironment": INSPIREseq Supplemental 8-2023

**Supplemental information**

**Supplementary Figures**

**Supplementary Figure 1. Enrichment of VHH across *in vivo* biopanning by percentage of unique VHHs. A.** Histogram showing the frequencies of CDR3 amino acid length for the library, Py117 lymph node (LN), and Py8119 LN samples. **B.** Percentage of unique VHHs in LN samples of Py117 and Py8119 tumor bearing mice. **C.** Distribution of unique CDR regions (CDR1, CDR2 and CDR3) showing the diversity is higher in CDR3 in comparison to other regions regardless of biopanning cycle.

**Supplementary Figure 2. Cell subpopulation enrichment using magnetic beads. A.** Scheme of immune cells sorting by magnetic beads. **B.** Flowcytometry gating strategy to select cells subpopulations CD11c$^{+}$, CD11b$^{+}$, CD8$^{+}$, CD4$^{+}$CD25$^{-}$, and CD4$^{+}$CD25$^{+}$ with fluorescence minus one (FMO) sample. **C.** Distribution of cells after Ficoll and dead cell removal. **D.** Enrichment of each cell type after magnetic beads separation.

**Supplementary Figure 3. Enrichment of VHH across samples of *in vivo* biopanning. A.** Rare clonal proportion showing summary proportion of VHHs with specific counts across biopanning in each cell type in Py117 and Py8119 tumor samples (upper panel) and Py117 and Py8119 LN samples (lower panel) from BP1 to BP4.

**Supplementary Figure 4. Diversity, rarefication, and clone tracking of *in vivo* biopanning in different tissues. A.** Rarefication curve assessed the diversity of BP1 to BP4 in Py117 and Py8119 LN samples through extrapolation and subsampling. **B.** True diversity or the effective number of types of BP1 to BP4 in Py117 and Py8119 tumor samples. **C.** Tracking Top VHHs in BP0 towards BP4 in Py8119 tumor (upper), Py8119 LN (middle), Py117 LN (lower) samples. **D.** Tracking Top VHHs in BP4 backward to BP0 Py8119 tumor (upper), Py8119 LN (middle), and Py117 LN (lower) samples.

**Supplementary Figure 5. Sanger and NGS details detected of VHHs from biopanning and scRNAseq sample assessments. A.** Total number of sequenced VHHs by Sanger sequencing and number of unique CDR3 sequences. **B.** Total number of detected VHHs by NGS sequencing and the number of unique CDR3 sequences. **C.** Total number of enriched VHHs in BP3 and BP4 and the number with unique CDR3 sequences. **D.** Overlap between detected VHHs by Sanger and NGS sequencing. **E.** Overlapping between detected VHHs in BP3 and BP4 by Sanger and NGS sequencing. **F.** Distribution of VHH clones detected by Sanger sequencing in NGS detected VHHs in BP0 to BP4.

**Supplementary Figure 6. Shard and overlapping nanobodies for enriched cells in the microenvironment. A.** Shared and unique VHH clones that bind to CD45 cells, enriched in biopanning 3 and 4, between tumor and lymph node in the Py117 and Py8119 models. Shared and unique clones for the (**B.**) CD8-VHH and the (**C.**) CD11c-VHH populations of enriched cells in each tissue and tumor.

**Supplementary Figure 7. scRNAseq sample assessment immunophenotype. A.** Single cell RNA sequencing data clustered by samples showing the harmonization and batch effect removal after integration. **B.** Cells distribution as proportion across all samples. **C.** Expression level of some canonical markers for cell sub population immunophenotyping of the scRNAseq. Mice were injected with PBS, inserted less phage, or libraries generated from sorted CD45, CD8, or CD11c cells from tumor bearing mice.

**Supplementary Figure 8**. **Nb1 antigen target identification. A.** Sliver stain of gels after immunoprecipitation of the Nb1 VHH with His-tag after incubation with splenocyte membrane extract lane 2 and no extract in lane 3. The band with the red arrow was observed after three independent preparations both methodologies and subsequently processed for identification. **B.** Mass Spectrometry results table of the most abundant proteins after gel extraction of the identified band.

**Supplementary Figure 9. Nb1 binds extracellular PHB2 on human cells. A.** Colocalization of Nb1 clone with PHB2 on H1299 human cells. Confocal image of fixed H1299 cells stained with α-PHB2 antibody and nanobody (upper panel) or stained with secondary α-VHH antibody and tertiary anti-mouse AF488 / anti rabbit-APC antibodies (lower panel) 24 h after seeding on coverslip. Höchst33342 nuclear stain (blue). **B.** Mander's colocalization coefficient M1 in H1299 cells, showing colocalization of α-PHB2 with Nb1 staining. The Nb1 channel pixel overlap with PHB2 channel (mean ± SE) for Nb1and α-PHB2 (n = 20 images), α-PHB2 antibody alone (n = 6) and 2nd/3rd antibodies alone (n = 6) from independent triplicates with six incrementing pixel shift configurations.

**Supplementary Figure 10. Structure prediction with Rosetta and AlphaFold A.** Structural model of the pentameric assembly of the coil domain (rainbow colors) and binding energy vs RMSD calculations with Rosetta. Each single monomer in the assembly strongly interacts with the remaining four monomers, suggesting that this conformation is extremely stable. **B.** AlphaFold prediction of the full structure of a monomer of PHB2. **C.** SnugDock binding energy vs RMSD results for the 11 docking sites. The cut-off of -40 REU (dashed line) or lower indicates the minimal requirement for binding. Only Sites 1, 5, 6 and 8 pass the cut-off with more than 10 models each. **D.** Structural clashes of Nb1 with the multimeric version of PHB2. Docking site 5 (left panel) and Docking site 6 (right panel) are not accessible when PHB2 is in a multimeric state (5x_CC model in gray).

### Supplementary Tables

Supplementary Table 1. Basic statistics of NGS detected VHHs in tumor and LN samples

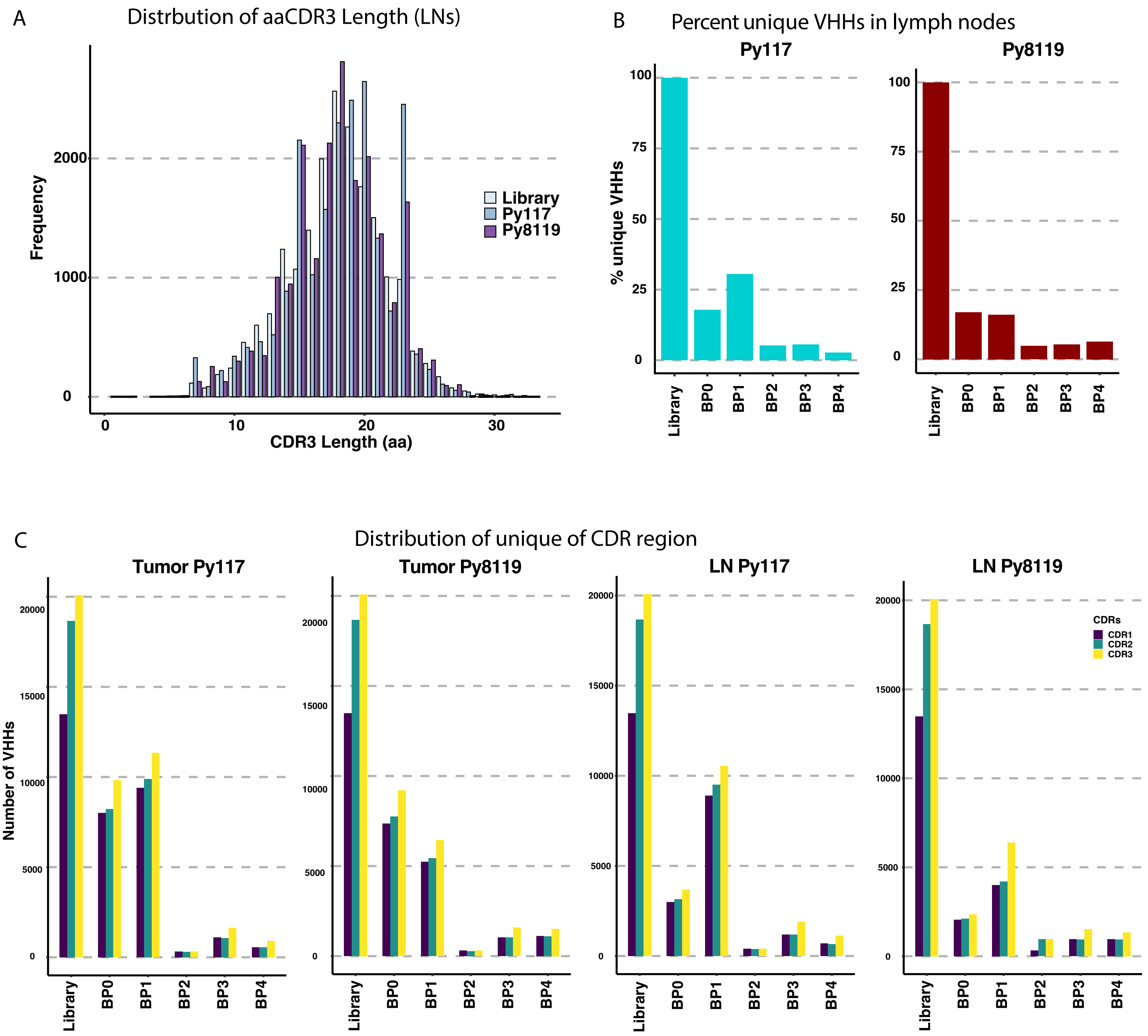


Supplementary Figure 1

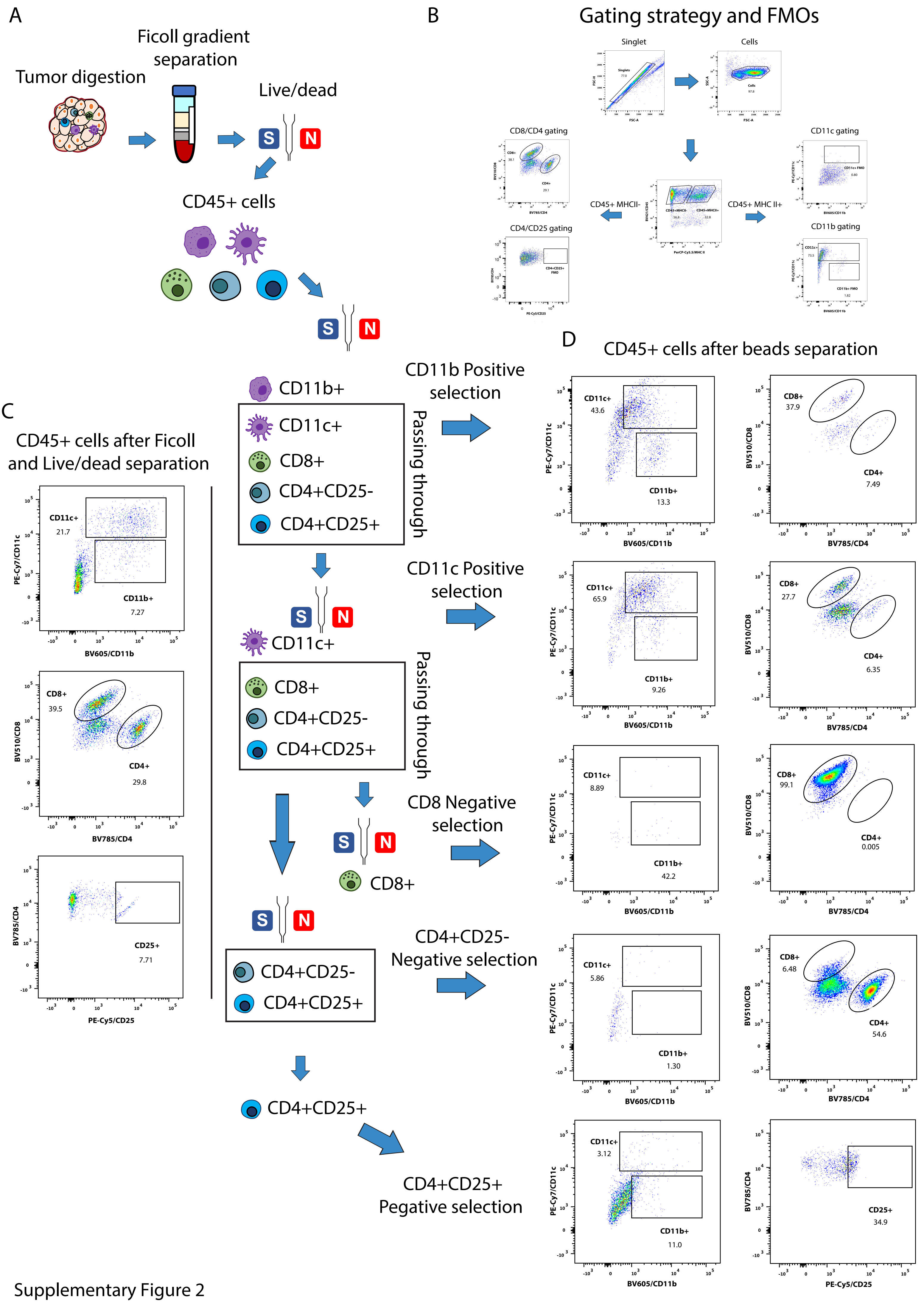


Supplementary Figure 2

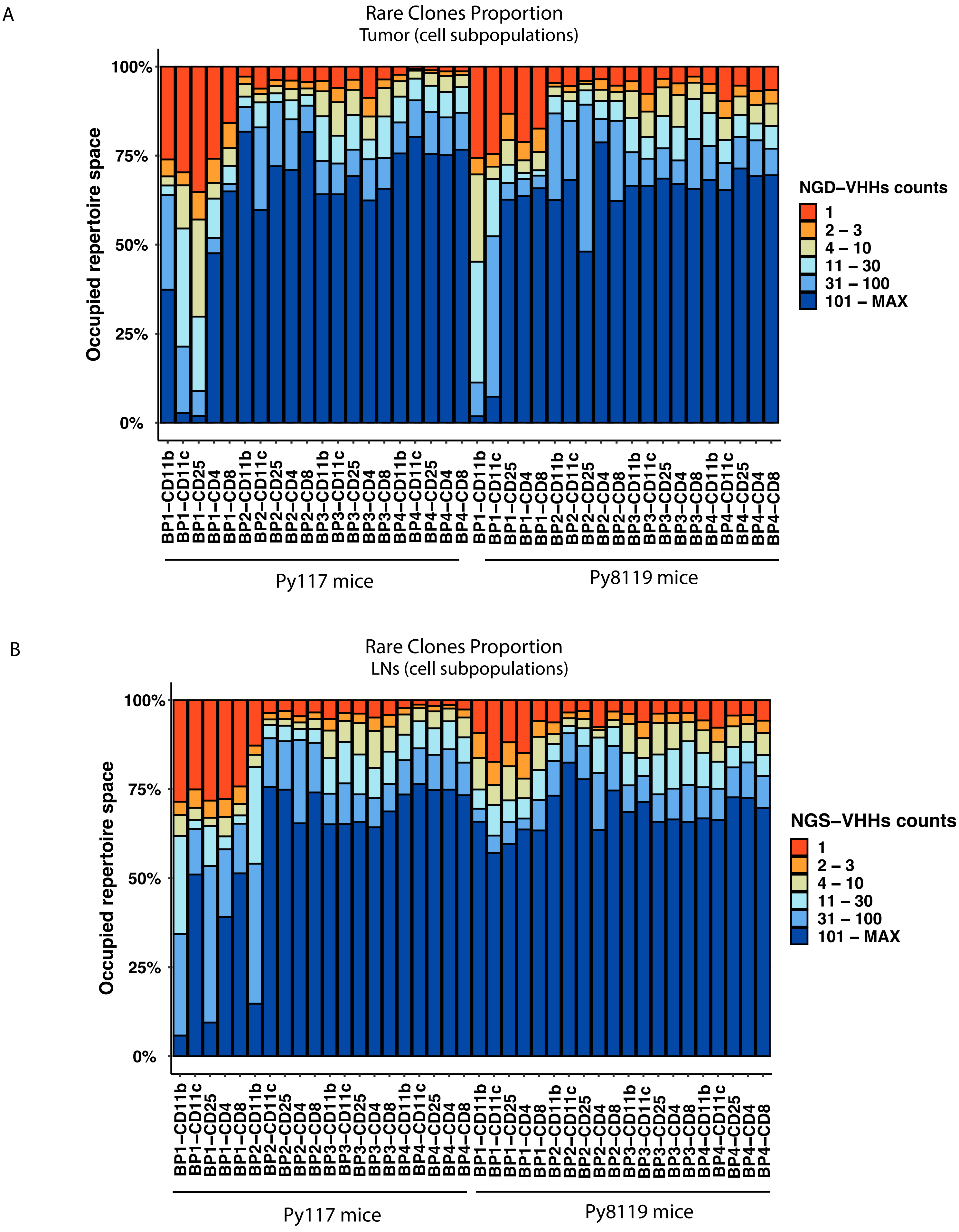


Supplementary Figure 3

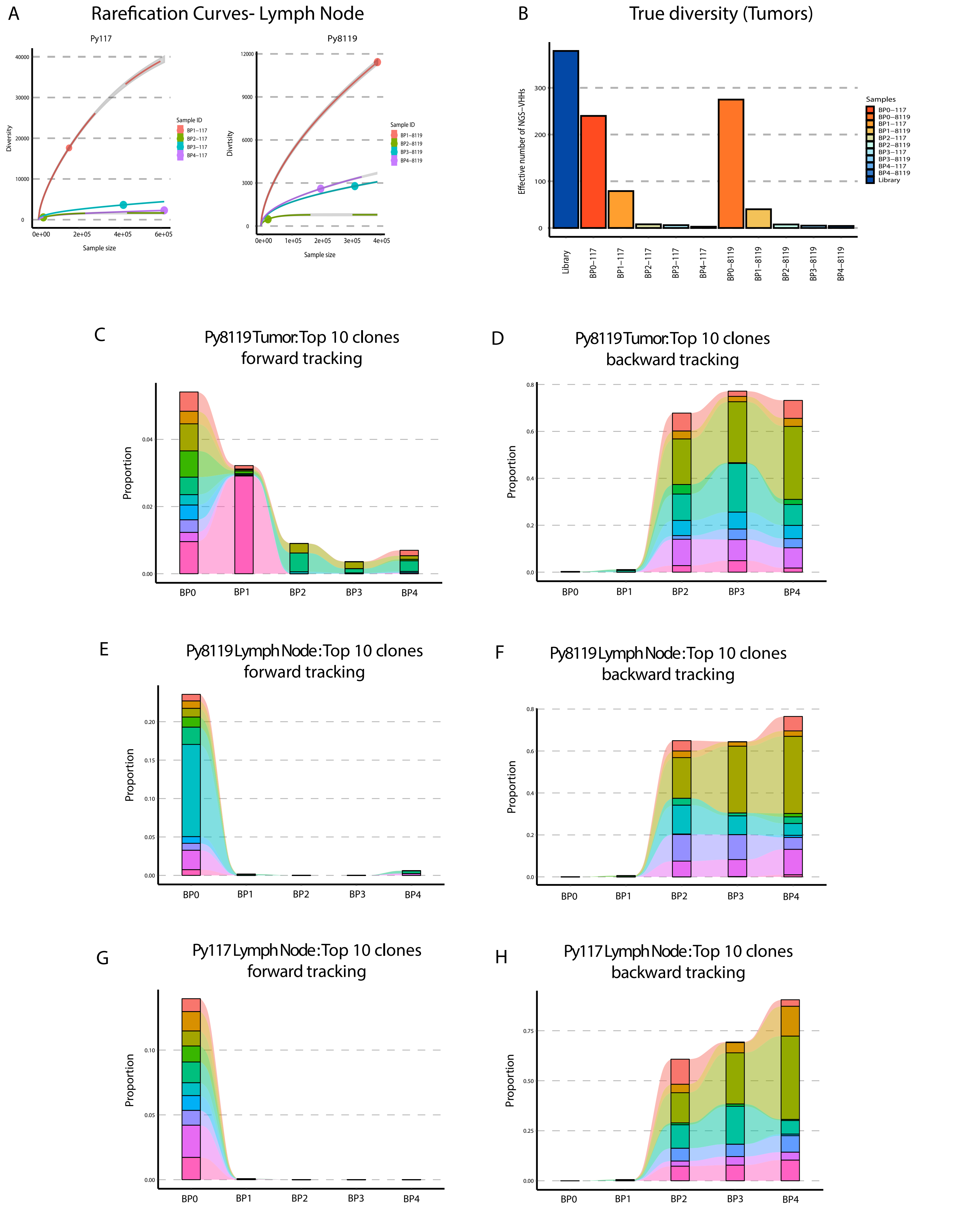


Supplementary Figure 4

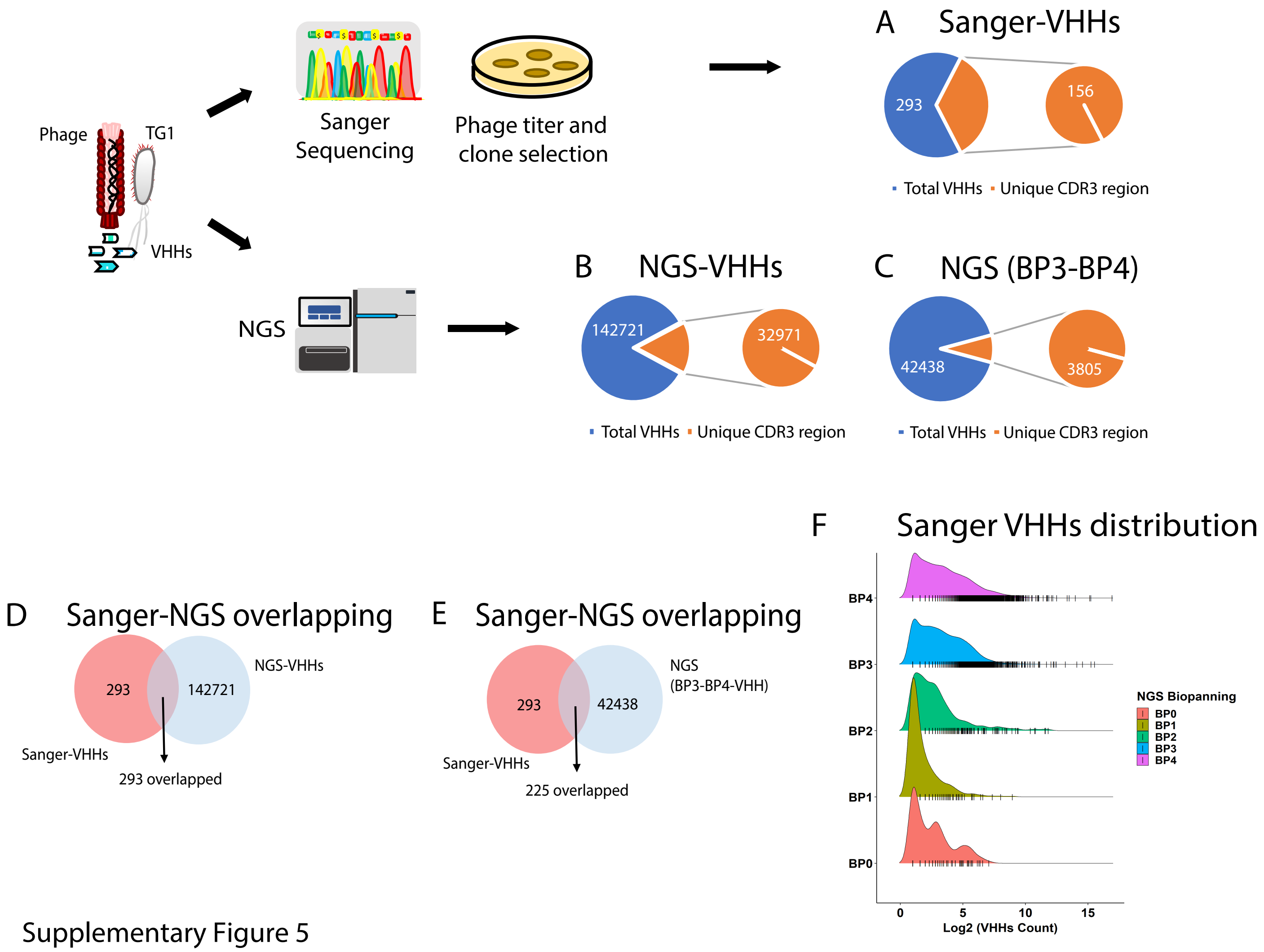


Supplementary Figure 5

A

### Shared and unique VHHs between tumor and LN in Py117 and Py8119

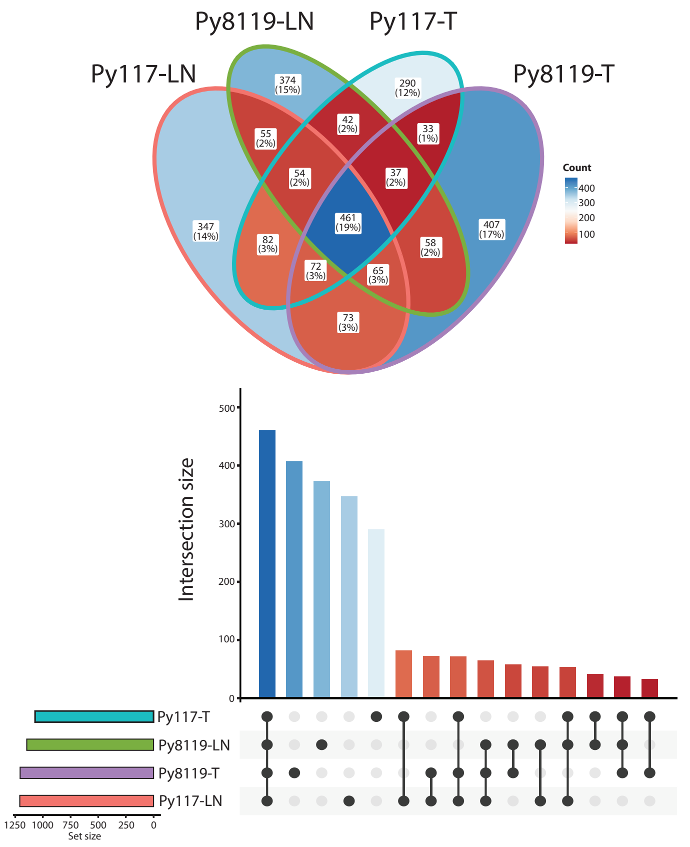


B

### Shared and unique CD8-VHHs between tumor and LN in Py117 and Py8119

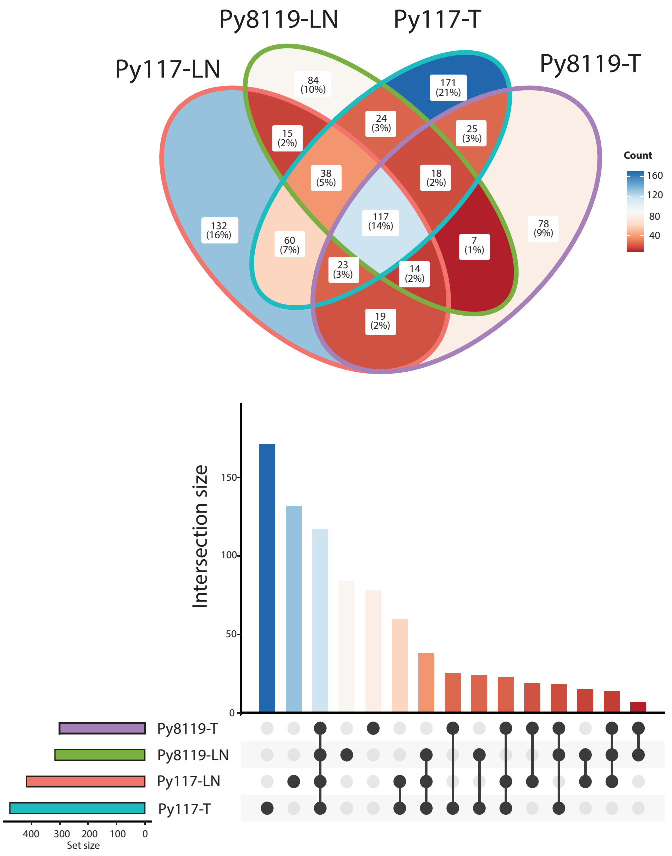


C

### Shared and unique CD11c-VHHs between tumor and LN in Py117 and Py8119

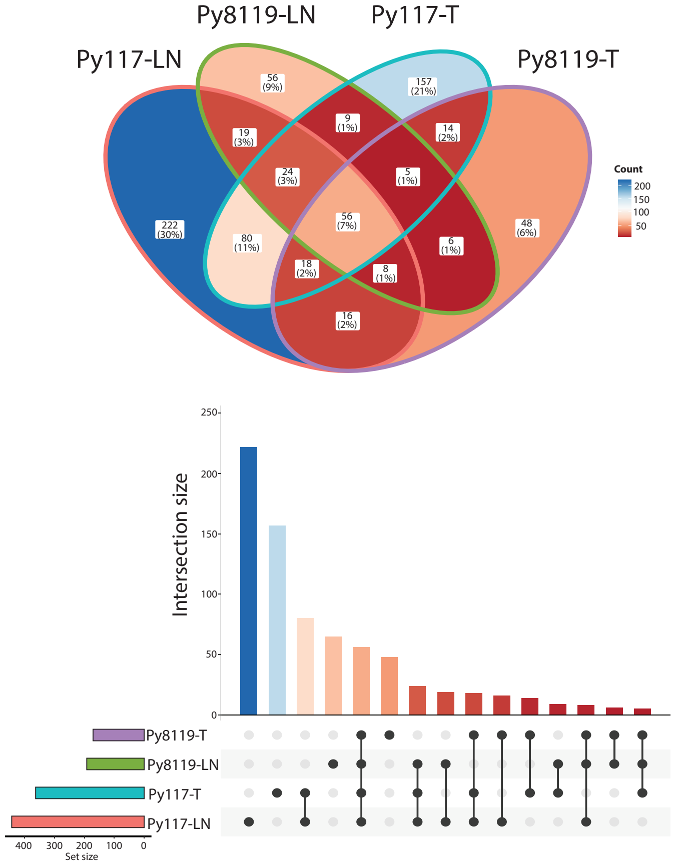


Supplementary Figure 6

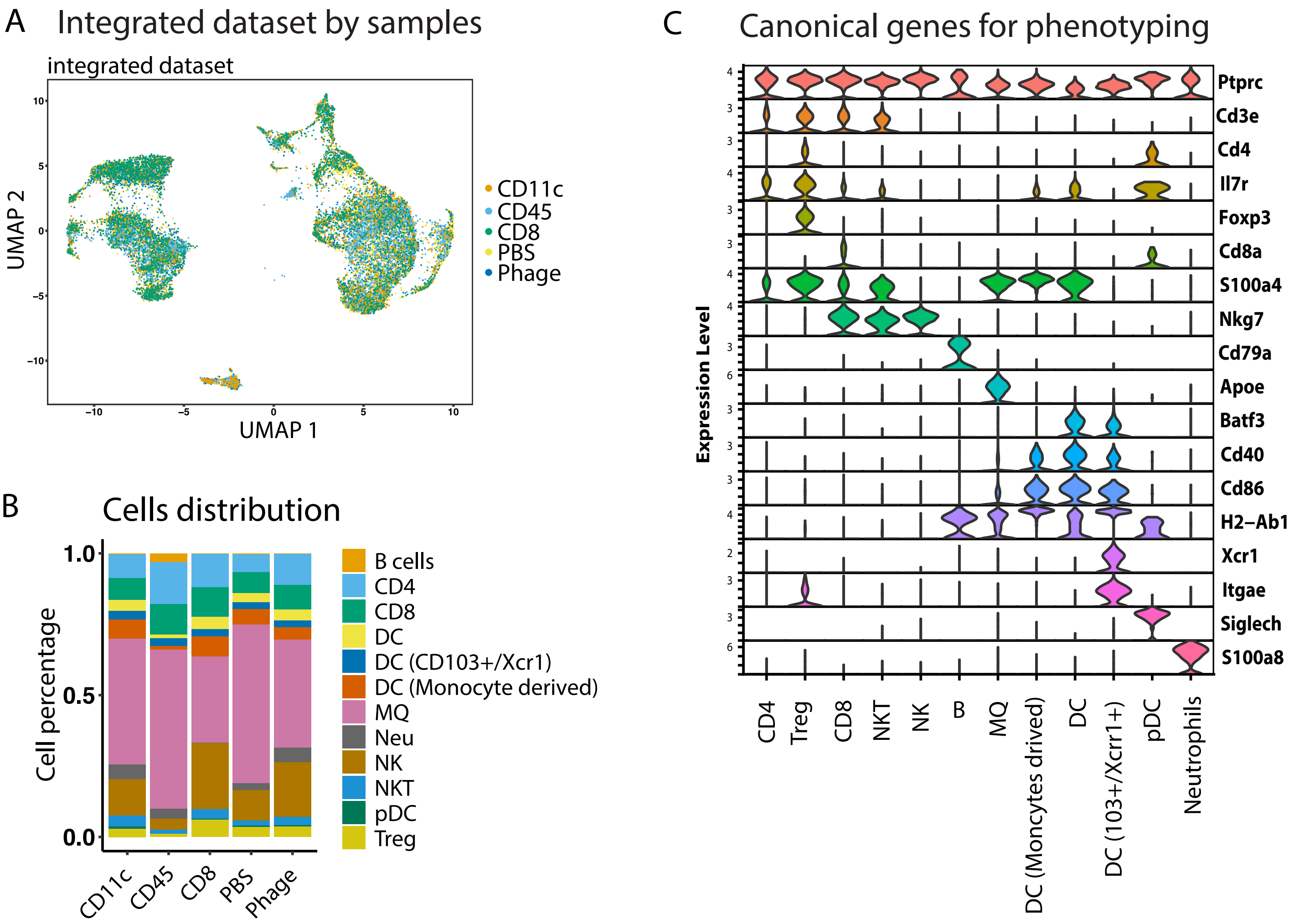


Supplementary Figure 7

A

Nb1 specific protein identification by Immunopreciptation

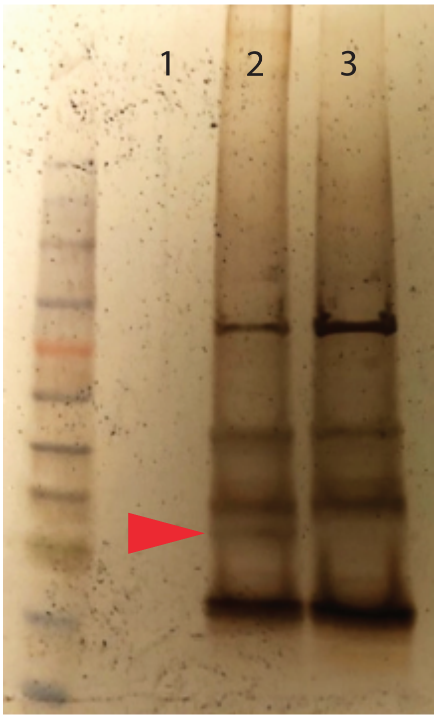


1- Dynabeads + Sp. MP

2- Dynabeads+Nb1-IgGFc+Sp.MP

3- Dynabeads+Nb1-IgGFc

B

Tandem MS of the Nb2 binding protein

| Accession | Description | MW in kDa | Gene Symbol | Abundance |
|---|---|---|---|---|
| O35129 | Prohibitin-2 | 33.3 | Phb2 | 1.72E+07 |
| P47911 | 60S ribosomal protein L6 | 33.5 | Rpl6 | 8.19E+06 |
| Q91WN1 | DnaJ homolog subfamily C member 9 | 30 | Dnajc9 | 3.43E+06 |
| G3UX26 | Outer mitochondrial membrane protein porin 2 | 30.4 | | 2.94E+06 |
| H3BKD0 | Heterogeneous nuclear ribonucleoprotein K (Fragment) | 33.2 | | 2.26E+06 |
| P14869 | 60S acidic ribosomal protein P0 | 34.2 | Rplp0 | 2.05E+06 |
| P14148 | 60S ribosomal protein L7 | 31.4 | Rpl7 | 1.55E+06 |

Supplementary Figure 8

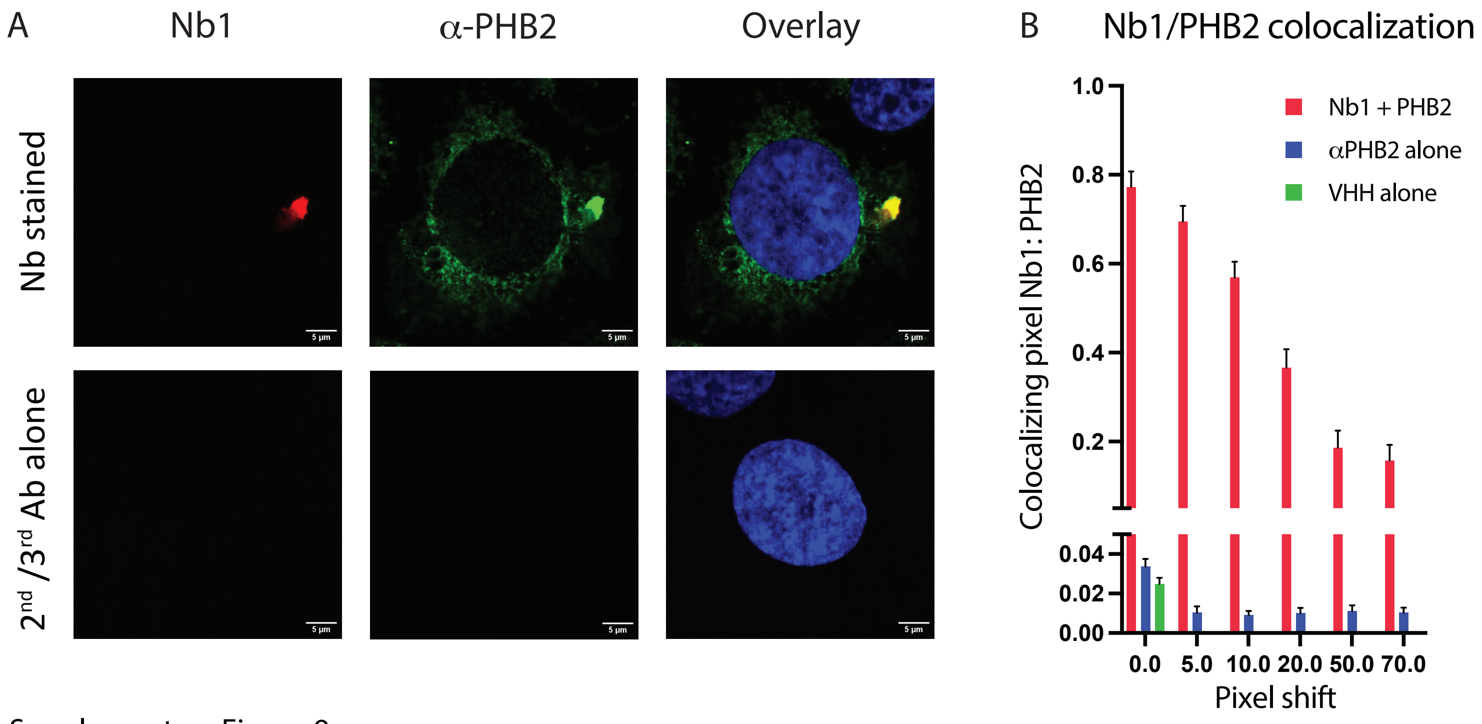


Supplementary Figure 9

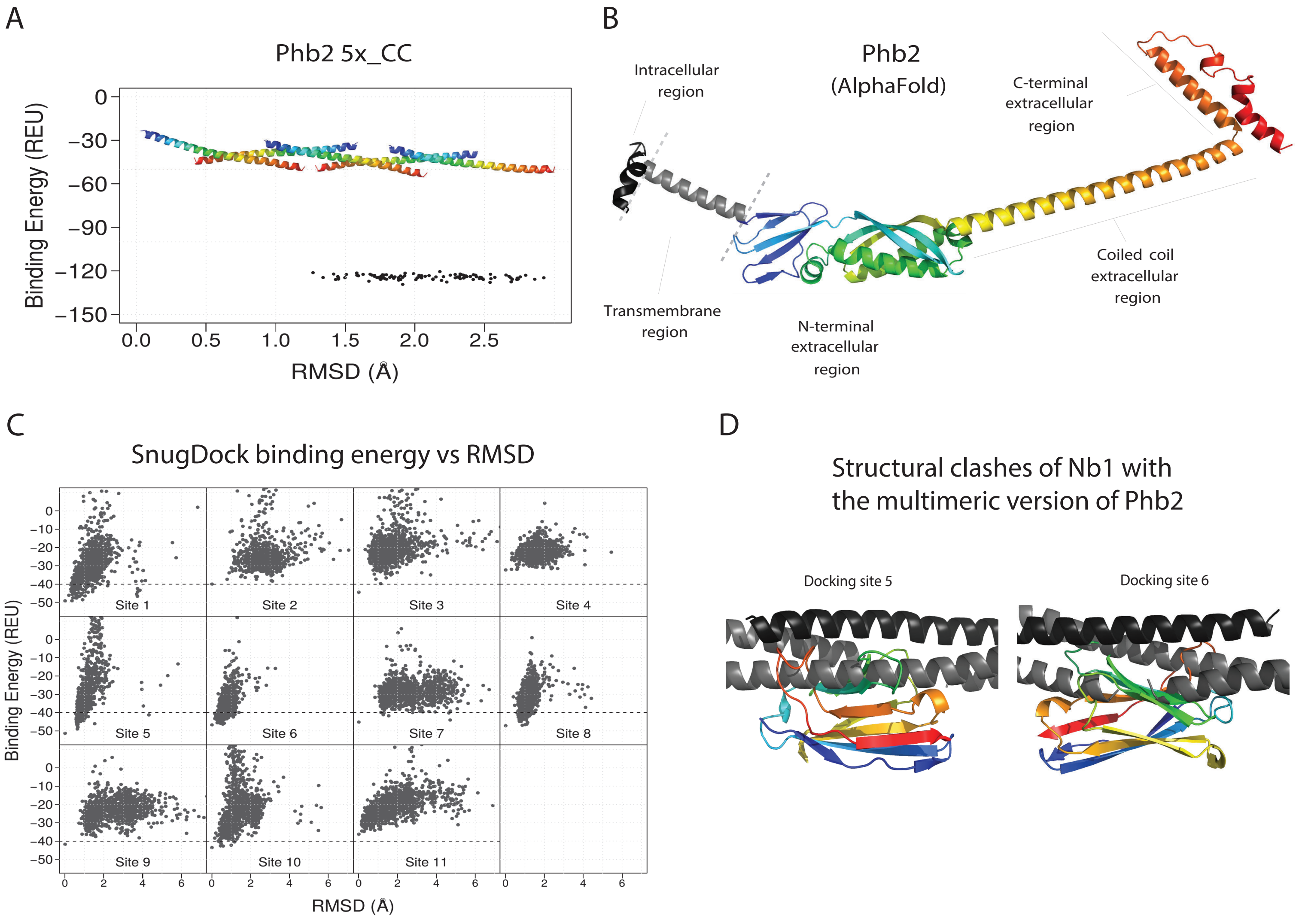


Supplementary Figure 10

Supplementary Table 1: Basic statistics of NGS detected VHHs in tumor and LN samples

| Biopanning | number of total assmbled VHHs | unique assembled VHHs | % of unique VHHs | Unique CDR1 | Unique CDR2 | Unique CDR3 |
|---|---|---|---|---|---|---|
| Library | 20083 | 20083 | 100 | 13476 | 18663 | 20083 |
| BP0-117-Tumor | 36095 | 15632 | 43.31 | 8013 | 8231 | 9833 |
| BP0-8119-Tumor | 21406 | 13593 | 63.5 | 7351 | 7754 | 9198 |
| BP1-117-Tumor | 164498 | 48383 | 29.41 | 9392 | 9896 | 11336 |
| BP1-8119-Tumor | 149591 | 33713 | 22.54 | 5232 | 5430 | 6430 |
| BP2-117-Tumor | 26475 | 1279 | 4.83 | 323 | 303 | 322 |
| BP2-8119-Tumor | 19599 | 1080 | 5.51 | 305 | 280 | 307 |
| BP3-117-Tumor | 314527 | 19431 | 6.18 | 1113 | 1074 | 1635 |
| BP3-8119-Tumor | 344630 | 17335 | 5.03 | 1030 | 1038 | 1576 |
| BP4-117-Tumor | 532758 | 11474 | 2.15 | 563 | 548 | 914 |
| BP4-8119-Tumor | 204625 | 15102 | 7.38 | 1121 | 1089 | 1515 |
| BP0-117-LN | 110950 | 19844 | 17.89 | 2996 | 3144 | 3678 |
| BP0-8119-LN | 61514 | 10442 | 16.97 | 2055 | 2110 | 2356 |
| BP1-117-LN | 149946 | 45703 | 30.48 | 8899 | 9505 | 10544 |
| BP1-8119-LN | 394966 | 63636 | 16.11 | 3998 | 4193 | 6393 |
| BP2-117-LN | 24104 | 1250 | 5.19 | 400 | 386 | 403 |
| BP2-8119-LN | 24467 | 1177 | 4.81 | 328 | 953 | 964 |
| BP3-117-LN | 416025 | 23468 | 5.64 | 1182 | 1191 | 1884 |
| BP3-8119-LN | 318519 | 17033 | 5.35 | 953 | 935 | 1519 |
| BP4-117-LN | 615924 | 16734 | 2.72 | 705 | 661 | 1133 |
| BP4-8119-LN | 202851 | 12914 | 6.37 | 964 | 942 | 1349 |
